# A simple technique to classify diffraction data from dynamic proteins according to individual polymorphs

**DOI:** 10.1101/2020.12.14.422680

**Authors:** Thu Nguyen, Kim L Phan, Dale F Kreitler, Lawrence C Andrews, Sandra B Gabelli, Dima Kozakov, Jean Jakoncic, Robert M Sweet, Alexei S Soares, Herbert J Bernstein

## Abstract

One often observes small but measurable differences in diffraction data measured from different crystals of a single protein. These differences might reflect structural differences in the protein and potentially reflect the natural dynamism of the molecule in solution. Partitioning these mixed-state data into single-state clusters is a critical step to extract information about the dynamic behavior of proteins from hundreds or thousands of single-crystal data sets. Mixed-state data can be obtained deliberately (through intentional perturbation) or inadvertently (while attempting to measure highly redundant single-crystal data). State changes may be expressed as changes in morphology, so that a subset of the polystates may be observed as polymorphs. After mixed-state data are deliberately or inadvertently measured, the challenge is to sort the data into clusters that may represent relevant biological polystates. Here we address this problem using a simple multi-factor clustering approach that classifies each data set using independent observables in order to assign each data set to the correct location in conformation space. We illustrate this method using two independent observables (unit cell constants and intensities) to cluster mixed-state data from chymotrypsinogen (ChTg) crystals. We observe that the data populate an arc of the reaction trajectory as ChTg is converted into chymotrypsin.

## 1. Introduction

Proteins often undergo structural changes as part of their normal functioning. Crystal structures often reveal proteins in different conformations (called polymorphs). Crystallography explores an average structure of all the molecules in the volume interrogated by the X-ray beam, often hundreds of thousands to hundreds of millions of molecules and possibly of significantly different polymorphs. These different structures might have revealed information on the dynamics of transitions among those polymorphs had they not all been averaged together. Because of that averaging, instead of seeing distinct states clearly, we may see only what looks like blurred thermal motion.

To reduce this problem, a typical structural study of a protein might involve somehow constraining the molecule in one of its states by binding an inhibitor before crystal growth. For example, in the emerging field of biological data storage, proteins with two distinct conformations (called polystates) are intentionally switched between them to represent binary code (0 and 1) (Sethi, 2015). It seems likely that molecules in a single crystal form would show slightly different structures depending on pH, the state of hydration, etc. That is, they exhibit dynamic behavior that may or may not indicate changes related to their function. Here we will use the term “polystates” to refer to protein structural polymorphs that correspond to biologically relevant conformations of proteins as well as to other significant state variations.

The normal course of modern crystallographic practice provides opportunities to discover and analyze the sort of changes we describe here. In particular, data collection at a synchrotron source often includes measurement of many partial sets of crystal diffraction data from many, often very small, crystals (Liu, Hendrickson, 2011) (Giordano *et al.*, 2012) (Rossmann, 2014) (Assman *et al.*, 2016) (Bernstein *et al.*, 2017) (Gao *et al.*, 2018) (Bernstein *et al.* 2020). This is possible because fourth generation synchrotron sources are very bright, the 2D detectors employed are very fast, and modern goniometers are very precise. A typical protocol takes about 9 seconds to do a full rotation of a 5 - 20 μm crystal by 0.2° for each image taken at 200 Hz. For this sort of treatment, one may mount crystals singly or, say, half a dozen in each sample mount (loop / mesh micro mount); in each case data are taken for each individual crystal. The crystals may come from different crystallization drops, or even different preparations, but they are all nominally isomorphous; our objective is to improve the quality of the data by a merging of multiple measurements. In fact, however, the data may be partitioned into different clusters of crystals, each cluster representing one step in the normal dynamic motion of the molecule. Starting from multiple independent samples increases the chances of having multiple states to observe.

There are two popular criteria for clustering datasets: similarity in lattice-parameter values or reflection intensities. Cell databases have long been used for substance identification and are now used as a coarse screen for molecular-replacement candidates. Steno (1669), as cited in Authier (2013), noticed the constancy of interfacial angles of crystals. Reflection intensities represent the true structure, so similarity of reflections is a good metric to use in comparing datasets.

To observe any such dynamic behavior – a different molecular state in each crystal – one must partition the single-crystal data on different measurable properties of each crystal. We demonstrate our approach using 146 diffraction-data sets that were obtained from single crystals of the protein chymotrypsinogen (ChTg) crystallized in three different crystallization conditions, pH 4.6, 5.6 and 6.5. Almost all of the crystals that were successfully assembled into solvable clusters were from crystals formed at pH 6.5. To detect different molecular states in individual crystals we partition the single-crystal data on these two properties: cell parameters and intensities. We first employed differences in cell parameters as a conventional method for clustering single crystal data. However, this was clearly inadequate, and we extended this to partitioning on similarities of diffraction intensities. Evidently a single perspective of an object fails to detect movements of that object which are invariant with respect to that single perspective. That is, we believe we have a clustering system that combines information from two or more observables to classify single crystal data more accurately according to the slightly different but stable conformational states that generated those data. In the end, as will be shown, in this case the clustering by correlation of intensities also reveals the large unit cell differences we observe, although it may be useful in some cases. In general. however, unit cell differences are observable much earlier in the course of structure solution than clusterable reflections and can be essential in doing sufficient preliminary clustering to be able to calculate meaningful correlation coefficients.

ChTg is the precursor (zymogen) for the mammalian digestive enzyme chymotrypsin (CHT), one of several well-known serine proteases (Kunitz, Northrop, 1935) (Siekevitz, Palade, 1960). This conversion is accomplished by several enzymatic cleavages. Firstly (in the digestive tract) trypsin cleaves the peptide bond between Arg15 and Ile16 to yield π-chymotrypsin, which is the active enzyme form. Secondly π-CHT molecules autolyze one another to cleave the bonds between Leu13 - Ser14 to release Ser14 – Arg15, and between Tyr146 - Thr147 and Asn148 - Ala149 to release Thr147 - Asn148. The resulting α-chymotrypsin is formed by three chains held by disulfide bridges.

## 2. Methods

### 2.1. Crystallization

The crystallization conditions for chymotrypsinogen (SIGMA) were determined using the commercial Hampton Crystal Screen HT (Hampton Research, Inc), and set up with a TTP mosquito robot (TTP Labtech/ STP Inc). Crystals were grown via hanging-drop vapor diffusion at 18°C from condition F11. To optimize the crystallization conditions further, we set up a 24-well tray with hanging-drop vapor diffusion with fixed pH of 6.5, varying both dioxane (10% or 15%) and ammonium sulfate (1.0 M-2.0 M). The drop, containing 1 μL of the reservoir solution (1.0 M-2.0 M ammonium sulfate, 0.1 M MES pH 6.5, and 10% or 15% dioxane) and 1 μL of 10.0 mg/mL of enzyme, was equilibrated over 0.5 mL of reservoir solution. The other two crystallization conditions had either 0.2 M ammonium acetate, 0.1 M sodium acetate trihydrate pH 4.6, and 30% w/v PEG 4000, or 0.5 M ammonium sulfate, 0.1 M sodium citrate, and tribasic dihydrate pH 5.6, with 1.0 M lithium sulfate monohydrate in the reservoir. All crystals were cryocooled in 3.5 M lithium sulfate. A typical drop of these crystals is shown in Fig. 1.

**Figure 1.**
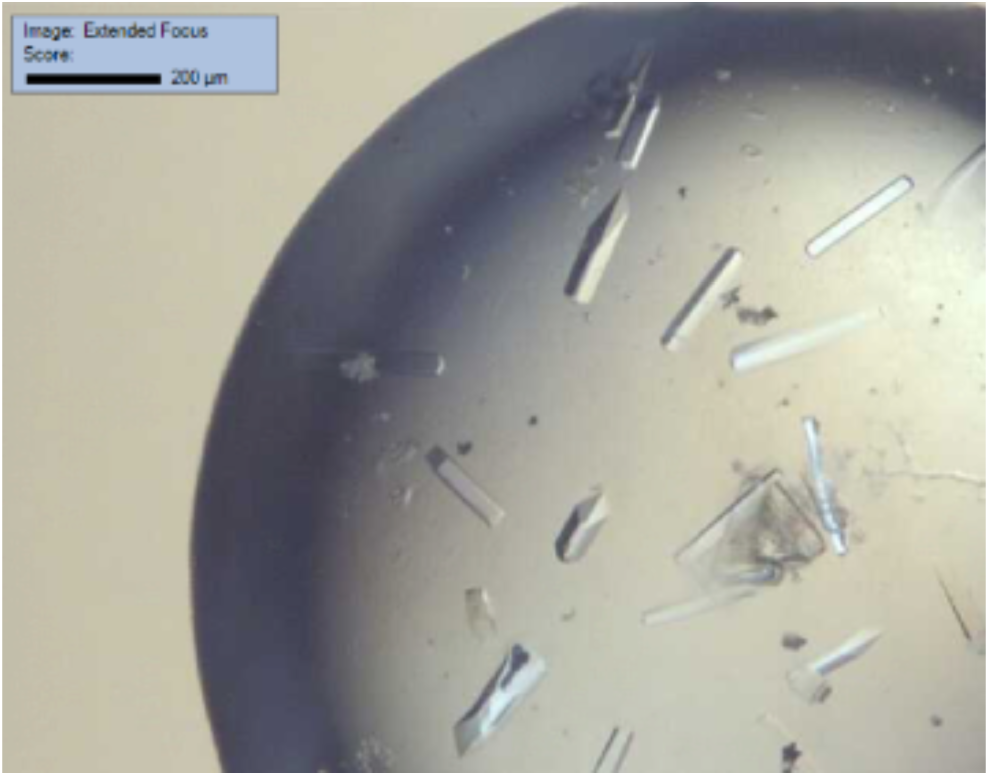
Representative ChTg crystals from the crystallization condition containing 1.0 – 2.0 M ammonium sulfate, 0.1 M MES pH 6.5, and 10% or 15% dioxane.

### 2.2. Data collection and structure determination

Data were collected at Brookhaven National Laboratory’s National Synchrotron Light Source-II at beamlines 17-ID-1(AMX) on a Dectris EIGER X 9M detector and 17-ID-2 (FMX) on a Dectris EIGER X 16M detector. The dataset for each crystal with sufficient data was indexed, integrated, and scaled using a version of fast_dp (Winter, McAuley, 2011) modified to run in the local distributed computing environment, XDS (Kabsch, 2010), DIALS (Winter *et al.*, 2018), PHENIX (Afonine *et al*. 2012), aimless and pointless (Evans, Murshudov, 2013). Tetragonal crystals of chymotrypsinogen diffracted between 2.0 - 2.4 Å. The structure was determined by molecular replacemement with PHASER (McCoy *et al.*, 2007) using ChTg PDB entry 1EX3 as a template. The data were refined to a final resolution using iterative rounds of refinement with REFMAC5 (Murshudov *et al.,* 1997) (Murshudov *et al.,* 2011) and manual rebuilding in COOT (Emsley, Cowtan, 2004). Structures were validated using Coot and PDB Deposition tools. The method yielded reasonable models for each of 145 clusters from which we identified 5 large but distinct clusters, each representing a distinct conformation. See Table 1. Figures were rendered in PyMOL (DeLano, 2002) and RasMol (Sayle, Milner-White, 1995) (Bernstein, 2000).

**Table 1.** Data collection and processing

|  | 7KTY<br>Average<br>structure<br>(clust. 145) | 7KU1<br>Green<br>structure<br>(clust. 139) | 7KU2<br>Red<br>structure<br>(clust. 140) | 7KU3<br>Cyan<br>structure<br>(clust. 141) | 7KTZ<br>Purple<br>structure<br>(clust. 131) | 7KU0<br>Yellow<br>structure<br>(clust. 138) |
| --- | --- | --- | --- | --- | --- | --- |
| Wavelength (Å) | 0.9201 | 0.9201 | 0.9201 | 0.9201 | 0.9201 | 0.9201 |
| Temperature (K) | 100 | 100 | 100 | 100 | 100 | 100 |
| Detector | Eiger 9M | Eiger 9M | Eiger 9M | Eiger 9M | Eiger 9M | Eiger 9M |
| Crystal-detector<br>distance (mm) | 172.65 | 177.07 | 175.87 | 172.20 | 171.20 | 171.91 |
| Rotation range per<br>image (°) | .2 | .2 | .2 | .2 | .2 | .2 |
| Total rotation range<br>(°) | 120 | 120 | 120 | 120 | 120 | 120 |
| Space group | P 41 21 2 | P 41 21 2 | P 41 21 2 | P 41 21 2 | P 41 21 2 | P 41 21 2 |
| a, b (Å) | 114.49 | 114.49 | 115.57 | 111.33 | 111.49 | 111.47 |
| c (Å) | 51.90 | 51.9 | 52.92 | 51.87 | 52.02 | 52.36 |
| $\alpha, \beta, \gamma$ (°) | 90.000<br>90.000<br>90.000 | 90.000<br>90.000<br>90.000 | 90.000<br>90.000<br>90.000 | 90.000<br>90.000<br>90.000 | 90.000<br>90.000<br>90.000 | 90.000<br>90.000<br>90.000 |
| Resolution range<br>(Å) | 2.00 -<br>29.69 | 2.39 -<br>29.69 | 2.19 -<br>28.89 | 2.00 -<br>29.13 | 2.00 -2<br>9.19 | 2.02 -<br>29.24 |
| Total No. of<br>reflections | 23850 | 14139 | 18927 | 22533 | 22696 | 22261 |
| Completeness (%) | 99.94% | 99.80% | 98.98% | 99.81% | 99.93% | 99.59% |
| I/$\sigma$(I) | 10.91 | 9.48 | 9.76 | 10.66 | 12.15 | 10.85 |
| Overall B factor<br>from Wilson plot | 53.96 | 61.87 | 53.31 | 33.72 | 29.15 | 30.16 |

**Table 2.** Structure solution and refinement

|  | 7KTY<br>Average<br>structure | 7KU1<br>Green<br>structure | 7KU2<br>Red<br>structure | 7KU3<br>Cyan<br>structure | 7KTZ<br>Purple<br>structure | 7KU0<br>Yellow<br>structure |
| --- | --- | --- | --- | --- | --- | --- |
| Resolution range (Å) | 2.00-29.69 | 2.39-29.69 | 2.19-28.89 | 2.00-29.13 | 2.00-29.19 | 2.02-29.24 |
| Completeness (%) | 99.94% | 99.80% | 98.98% | 99.81% | 99.93% | 99.59% |
| No. of reflections | 23850 | 14138 | 18926 | 22533 | 22696 | 22261 |
| Final $R_{\text{cryst}}$ | 19.13% | 22.08% | 20.97% | 18.15% | 16.18% | 16.74% |
| Final $R_{\text{free}}$ | 20.19% | 26.37% | 23.42% | 21.01% | 19.01% | 19.41% |
| No. of non-H atoms |  |  |  |  |  |  |
| Protein | 1786 | 1771 | 1778 | 1794 | 1786 | 1786 |
| Ligand | 15 | 5 | 5 | 15 | 15 | 15 |
| Water | 99 | 10 | 44 | 195 | 249 | 243 |
| Total | 1900 | 1786 | 1827 | 2004 | 2050 | 2044 |
| R.m.s. deviations |  |  |  |  |  |  |
| Bonds (Å) | 0.01 | 0.01 | 0.01 | 0.01 | 0.01 | 0.01 |
| Angles (°) | 0.86 | 1.00 | 0.86 | 0.77 | 0.80 | 0.74 |
| Average $B$ factors (Å<sup>2</sup>) | | | | | | |
| Protein | 57.93 | 67.43 | 58.69 | 36.60 | 28.11 | 28.83 |
| Ligand | 62.96 | 83.90 | 75.68 | 40.69 | 40.69 | 34.83 |
| Water | 58.66 | 63.04 | 56.13 | 42.32 | 37.08 | 36.80 |
| Ramachandran plot<br>Procheckccp4 |  |  |  |  |  |  |
| Most favoured (%) | 97.47% | 96.20% | 97.06% | 97.47% | 98.31% | 98.73 % |
| Allowed (%) | 1.69% | 2.95% | 2.52% | 2.11% | 1.27 % | 0.84% |

We used these crystals to obtain several hundred datasets at native energy (13.475 keV, 0.92 Å) and several hundred additional datasets at low energy (7.0 keV, 1.77 Å). The data yielded molecular-replacement solutions against the model from the PDB (Bernstein *et al.*, 1977) (Berman *et al*., 2000) ChTg entry 1EX3 (Pjura *et al.*, 2000). We then used this model to build our “average” structure from an average of all subsequent data sets at the “native” energy.

During the scaling together of data from the multiple crystals we had already noted clusters in the data. We suspected they originated from dynamic behavior in the ChTg crystals; that is, that each crystal carried just one average polymorph. We describe the clusters observed in the ChTg data.

### 2.3. Data-clustering program

We used a custom-modified version of the clustering pipeline KAMO (Yamashita *et al.*, 2018), which uses the clustering program Blend (Foadi *et al.*, 2013) to generate a dendrogram of the data sets. We expanded KAMO and Blend to allow two-factor clustering, so that initial “coarse” clusters are obtained by a partitioning of data sets into groups according to the similarity of their crystallographic unit cells, and then “fine” clusters are generated by further partitioning of each cluster according to the similarity of the amplitude data. (The modified software is available in github.com/nsls-ii-mx/blend and github.com/nsls-ii-mx/yamtbx.) We used this approach because determining a similarity score requires that two data sets have a large number of measured amplitudes in common, *i.e.* that there be a reasonably complete set of structure factors. For simple Pearson Correlation Coefficient (CC) calculations 70% completeness is needed. If a penalty is introduced for unmatched structure factors, completeness as low as 20 - 40% can be used (Bernstein *et al*., 2017) and we are studying the effect of even lower completeness for applying that method to partial data sets. Figs. 4 – 8 of the results section illustrate our clustering approach.

Unit cell parameters and amplitudes contain independent information that, combined, should result in improved clusters. For example, one expects differences in cell parameters to be sensitive to changes in the outer shape of the structure including changes in the presence or shape of ligands complexed at the surface of the protein, while differences in amplitudes would be sensitive to ALL conformational changes in the protein. Our approach was to combine the strengths of both sources of information sequentially to generate optimal clusters, although later we showed that in this case **the intensities alone discriminate between the two clusters based on the length of a**.

Since with space group P41212 cell parameters **a** = **b** and all cell angles are 90 degrees, there were only two free parameters, so we can visually demonstrate the lattice clusters in terms of just cell parameters **a** and **c** for each dataset. We plotted the two values to demonstrate clusters (Fig. 4). Then we built the structure for each cluster based on the average structure, employing the merged data for each cluster.

Secondly, given a particular lattice, we wanted to demonstrate the changes of the structures based on structure factors. We used KAMO to cluster the datasets into different groups based on intensity CC. The distance between datasets is calculated by *d*(*i*, *j*) = √(1 − *CC*(*i*, *j*)). The method then uses a hierarchical clustering analysis (Rokach, Maimon, 2005) with Ward’s method (Ward, 1963) to form distinct groups of datasets. In Ward clustering, the datasets are considered first by building a small cluster out of the two closest datasets and then by adding one dataset at a time to whichever dataset or existing cluster results in a new cluster of smallest variance. There are many other choices of what is called “linkage” in forming a cluster dendrogram, such as using cluster centroids, but using the minimal variance allows use of one simple distance matrix as input to the clustering algorithm, rather than requiring repeated calculation of distances among cells, or, worse, among *hkl-*vectors of structure factors, but it does tend to produce dendrograms for which the heights grow rapidly. See (Strauss, von Maltitz, 2017) for some alternative linkage choices. The program outputs a dendrogram that illustrates the distances (differences) among clusters by the y axis (the height). To get a certain number of clusters which contain more similar datasets, we choose a height cutoff value k accordingly. The lower the k value is, the more similar the datasets in each cluster are. Each cluster now relates to a structure built after merging datasets within it.

We employed the average structure defined in Section 2.2 as the starting model for structure determination and refinement for each of the clusters’ structures. All the processes we used to build clusters’ structures and refine them later are automated with the help of REFMAC. Following the automated refinement steps, we performed a manual check-and-refine step using COOT to ensure no serious errors remained from the automated process and corrected the refined model as needed. FATCAT allowed us to quantify the morphological differences among structure solutions.

After determining the cell-parameter clusters and structure-factor clusters, we aligned the datasets of the latest clustering results with the cell parameter results to observe any relationships among calculated results from the same data.

### 2.4. Illustrating the differences to identify physically meaningful clusters

Any software that uses observable parameters to generate clusters may generate a very large number of clusters. How is one to determine which clusters are physically meaningful? Dendrograms can illustrate the relationships among clusters, but one must illustrate physical relevance using structural tools, *i.e.* comparing the structures obtained from each of these clusters. We generated two software tools for this purpose (see https://github.com/nsls-ii-mx/chymotrypsinogen). Both tools use individual colors to differentiate among clusters, which we can then test for physical relevance, and both tools use two- or three-dimensional plots to illustrate an underlying physical characteristic of the structure.

We developed a tool to create color-coded coordinate ellipses. We plotted the XYZ coordinates for the C_α_ atom of a particular amino acid in the structure that we observed to be highly mobile among the clusters. We created color-coded ellipsoids that enclosed all of the C_α_ atoms that we found to come from each of the individual clusters. The size of each ellipsoid indicates the variation of the coordinates within the corresponding cluster. Ideally the size of each color-coded ellipsoid will not be very large compared to the separations among the centroids of the ellipsoids, indicating that each cluster represents a separable state. The code is available in the github.com/nsls-ii-mx/chymotrypsinogen.git git repository in the file raw.githubusercontent.com/nsls-ii-mx/chymotrypsinogen/master/ellipsoid.py.

We also plotted the **a** and **c** axis lengths for each dataset that resides within an amplitude-based cluster in Fig. 4. We illustrated all data that originated from each postulated cluster in a different color in Fig. 6.

To detect subtle differences among the clusters’ structures, we used FTMap (Kozakov et al., 2015), software designed to determine and characterize ligand-binding hot spots on proteins’ surfaces. The algorithm uses a library of 16 molecules as probes to discover potential patches on the surface of a structure where a molecule might bind. Differences in proposed surface binding could reveal otherwise unnoticeable physical differences among the structures.

## 3. Results

### 3.1. Data collection and protein structures

We obtained several hundred native data sets (NSLS II 17-ID-1 and 17-ID-2, see Table 1) of which 146 could be merged. The protein is a single chain of 245 residues, of which four residues (147 – 150) are not resolved.

We obtained our initial structure, PDB accession code 7KTY, from a merge of all 146 datasets; we choose to call this the Average Structure (denoted thus in Table 1). We used REFMAC and COOT to refine the structure and reduce the R value to about 18%.

Averaging all 146 data sets together resulted in a relatively high Rmerge value (48%) but nevertheless 7KTY was a good fit to these data (Rwork 19%, Rfree 20%. This average structure is slightly different from the published 1EX3 structure which we used as an initial phasing model. For example, 7KTY has a missing loop from residue 147 to residue 150.

We show the comparison between the sequences for the 1EX3 ChTg structure and for our structure in Fig. 2.

**Figure 2.**
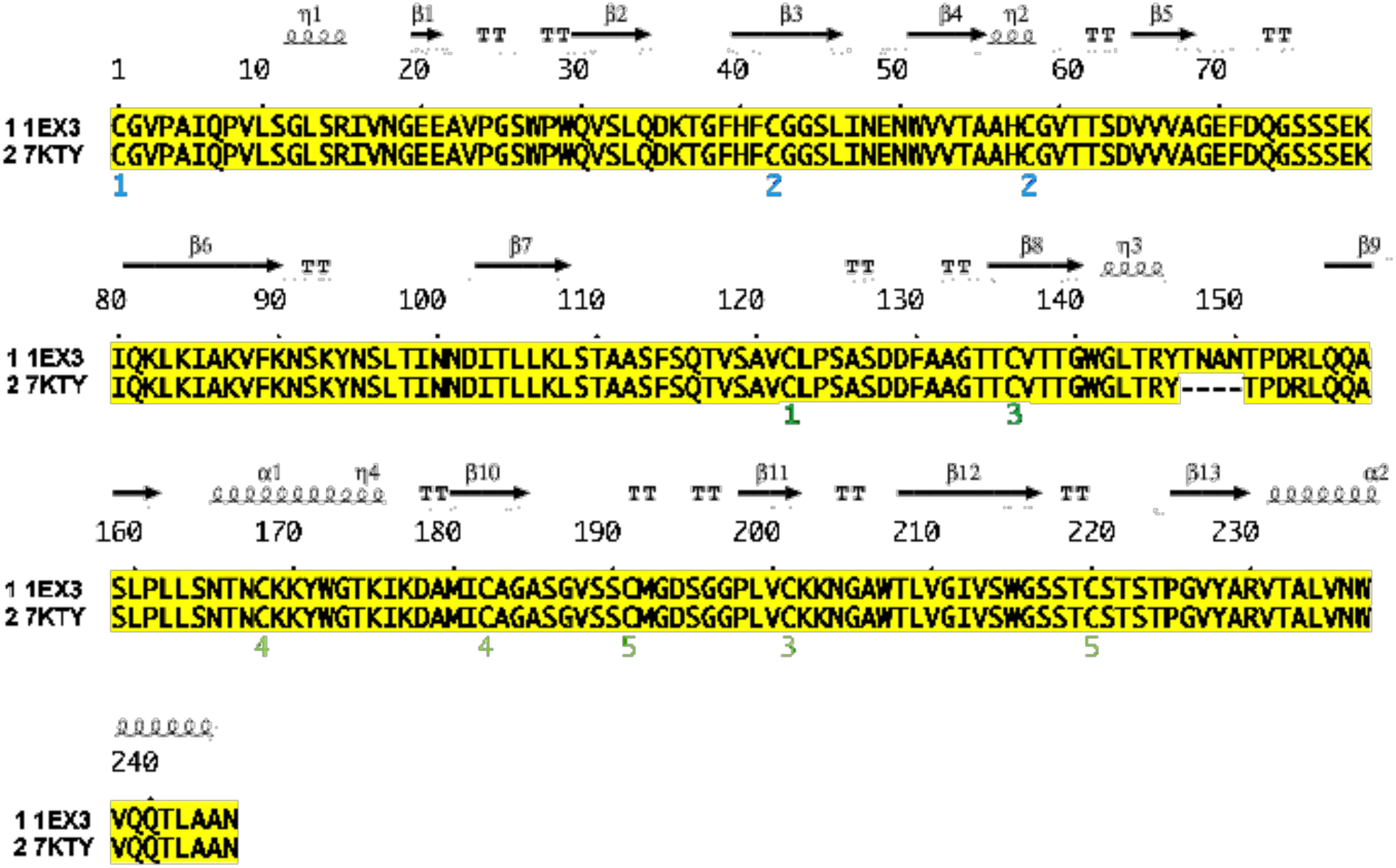
Sequence differences between chymotrypsinogen 1EX3 and our average structure 7KTY. We compare the two sequences, one from the PDB 1EX3 structure and one from our average structure, 7KTY. The figure shows that our structure has a missing loop from residue 147 to 150.

**Figure 3.**
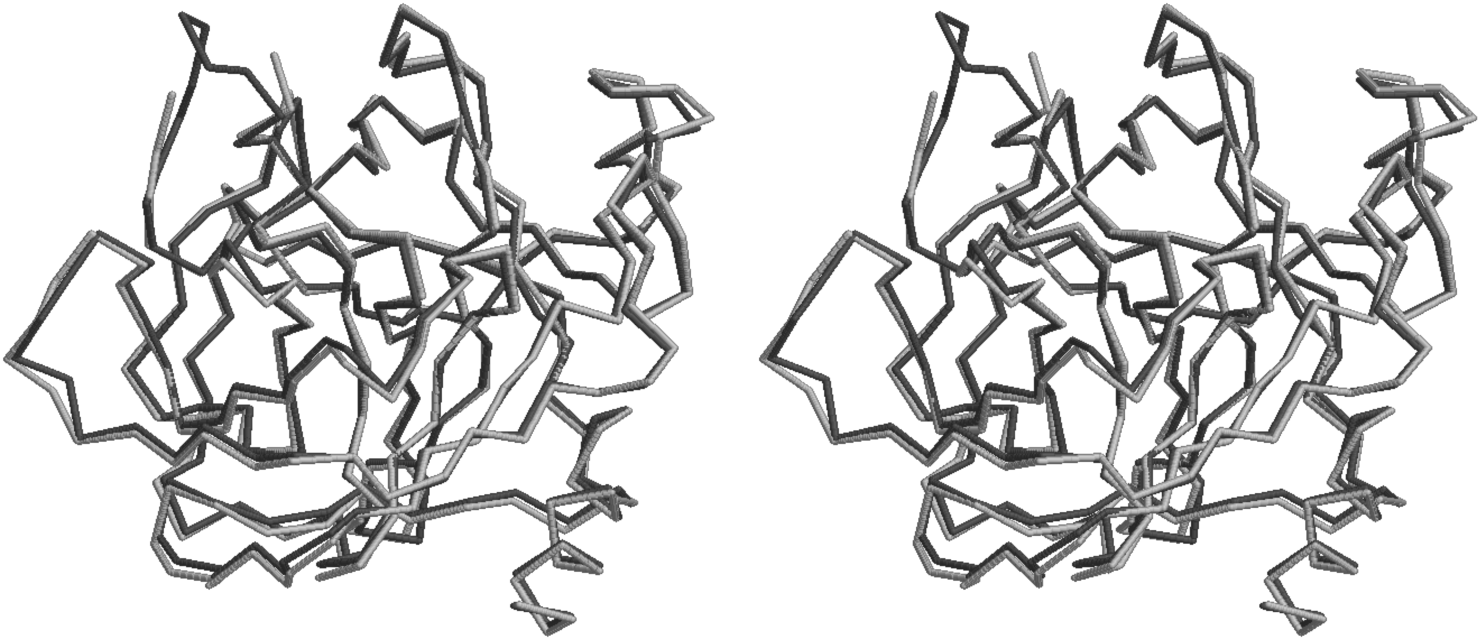
The published 1EX3 structure compared with the average structure 7KTY in cross-eyed stereo. We visually compare the PDB 1EX3 structure in dark grey and our average structure 7KTY obtained after merging all of our 146 data sets in light grey. The FATCAT chain RMSD is 0.56 Å. There were two regions with significant differences; they are adjacent, and appear at the upper left of this figure. First, in the average structure the amino acids between Thr147 and Asn150 are missing. Second, in the average structure the amino acids between Thr139 and Tyr146 adopt a significantly different conformation.

### 3.2. Clustering with unit cells and with amplitudes

Clustering software will generate data corresponding to candidate polystates, even in cases where truly distinct polystates are not actually present in the samples. Two independent data sets collected on two samples will always give different average structures. Such differences often are not relevant in terms of dynamics or states when the differences are small compared to experimental error. The only way to determine if candidate clusters may correspond to biologically relevant polystates is to generate and examine corresponding structural models (typically atomic models) with appropriate real-space tools, such as FATCAT and COOT. In the case of the ChTg data, we could see from inspection that the data could divide into two large clusters corresponding to structures with **a** ≈ 111 Å axis and those with **a** ≈ 114 Å. See Fig. 4.

**Figure 4.**
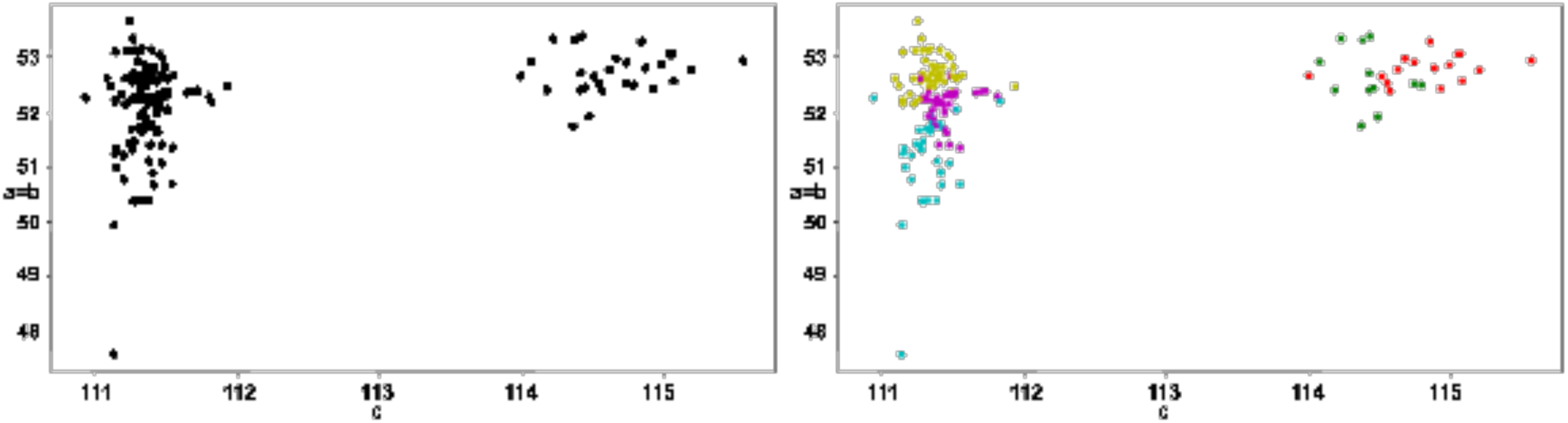
Two main data clusters can be identified by inspection (111Å group and 114Å group). We observed that our data partitioned cleanly between 28 data sets with an **a** (=**b**) unit cell of approximately 114Å and 118 data sets with an **a** (=**b**) unit cell of approximately 111Å. The separation into the two unit-cell clusters is shown in the monochrome clustering on the left. The further division of those two clusters into amplitude-based clusters is shown by the colors on the right. The 114Å A-cell-based cluster contained the green and red clusters, and the 111Å A-cell-based cluster contained the cyan, purple and yellow clusters. Each of our data sets was sufficiently large that amplitude-based clustering could have been used from the start. However, many serial crystallography projects consist of narrow wedges of data, each of which might be too small to cluster effectively using amplitudes because amplitude-based clustering requires that data sets have a sufficient number of observations in common. This figure illustrates how cell-based clustering might be used to boot-strap amplitude-based clustering (cell-based clustering would be used first in such cases, followed by amplitude-based clustering).

Employing only the observed diffraction intensities, we identified two main clusters that corresponded to the two main polymorphs that ChTg adopted in our crystals, based on the length of the **a** axis. In addition, there were five clusters that corresponded to biologically relevant polymorphs present in our data. The cell-based clustering shows that the **a** cell lengths separate clearly into two groups, while the **c** cell length varies less and is not clearly separable. There were significant solvent-region differences between the 111 Å **a** cluster and the 114 Å **a** cluster (Fig. 4).

When comparing the structures corresponding to the 111Å **a** cluster and the 114Å **a** cluster we observed that the 114 Å **a** cluster data yields observable density for all 245 residues (similar to 1EX3), while the 111 Å **a** cluster data indicates that there is a missing loop from residue 147 to residue 150 (this region is also not observed in the average structure). Another thing we observed is the presence of strong density near Lys 175 in the 111 Å **a** cluster data while the 114 Å **a** cluster data does not have this large artifact. See Fig. 5.

**Figure 5.**
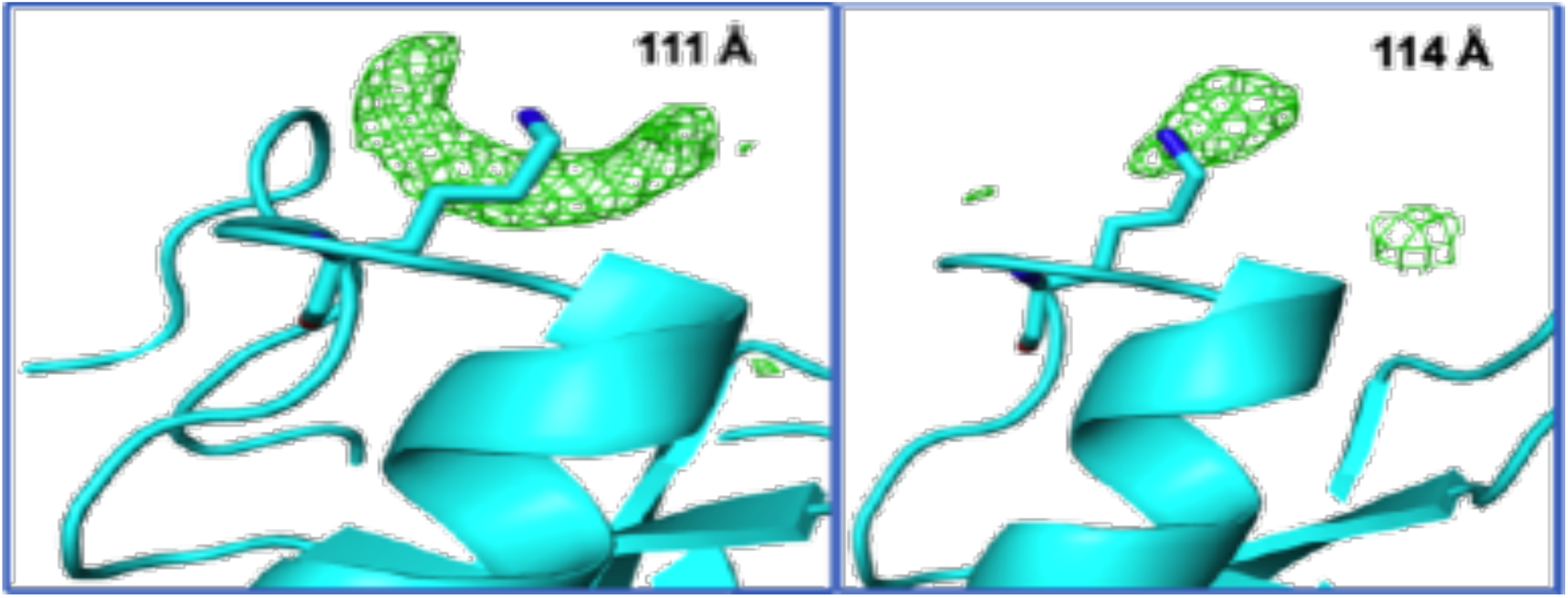
Differences in solvent between the **a** = 111 Å cluster and the **a** = 114 Å cluster. The electron density outside the protein envelope for the **a** = 111 Å cluster contains a semi-circular arc of density around the Lys175 nitrogen, while the corresponding region in the **a** = 114 Å cluster electron density contains a distinct density peak that we modelled as a water molecule.

Using Blend and KAMO, we obtained 145 clusters from the 146 ChTg data sets. We then generated structures after merging datasets belonging to each of these 145 clusters, and we visually inspected each of them to find any recurring patterns. This visual inspection allowed us to determine that all the reproducible differences could be accounted for by using just five of the larger clusters (which we call the green, red, cyan, purple and yellow clusters). In other words, we chose the “height” at which we cut the KAMO dendrogram so that five clusters contain the data corresponding to the relevant structures (Fig. 6). The 145 clusters could also be overlaid on the **a** and **c** axis diagram, color coded according to each of the five main clusters (Fig. 4). All the datasets of the green and red clusters belong to the 114 Å **a** cell-based cluster and datasets of the cyan, purple and yellow clusters belong to the 111 Å **a** cell-based cluster. If we increase the cut height to 1.5, we get the two intensities’ sub-master clusters, one containing green and red clusters, the other containing cyan, purple, and yellow clusters. This means the intensity cluster result has a strong alignment with the cell parameter cluster result.

**Figure 6.**
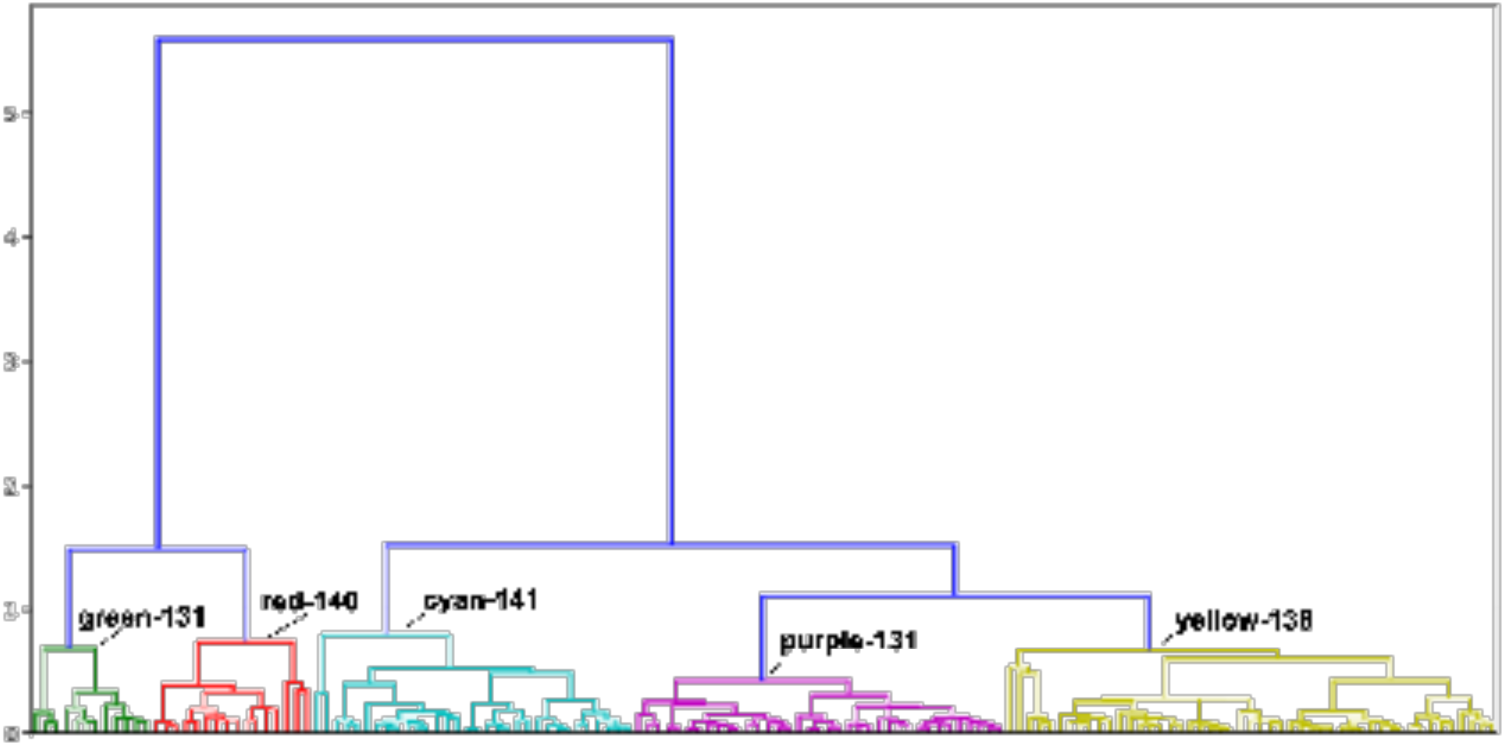
Amplitude-based clusters generated using KAMO (dendrogram). This dendrogram shows a representation of the similarity of pairs of data sets and clusters of more data sets. They are arranged with the most similar near one another, and the connecting bar at a height corresponding to the distance between clusters. The difference was calculated using Ward’s method for hierarchical clustering, which yields a composite metric that contains information from amplitude differences and from unit-cell differences. Our algorithm is described in Section 2.3. Structures were solved corresponding to all of these 145 clusters. We deposited the overall average structure as PDB entry 7KTY, and five distinct structures each corresponding to the highest level of each of the green clusters as PDB entry 7KU1, the red clusters as PDB entry 7KU2, the cyan clusters as PDB entry 7KU3, the purple clusters as PDB entry 7KTZ, and the yellow clusters as PDB entry 7KU0. We selected a height within the dendrogram at which to partition our data. We made our final choice to use five clusters through inspection of the derived structures.

### 3.3. The five data clusters

We generated a dendrogram using Ward’s method for hierarchical clustering with the height cutoff at 1.0 to get distinct groups of datasets. To observe the differences between the 145 structures generated using individual data clusters, we calculated the largest differences in physical coordinates at each residue C_α_. We observed that the most mobile area is near the missing loop from residue 146 to residue 152 (particularly residue 139 to residue 145). The largest differences were observed for residue 146 (with more than 3 Å average positional differences; Fig. 8).

At each residue position, we plotted ellipsoids to illustrate the variation in the C_α_ coordinates observed in each of the structures corresponding to the green, red, cyan, purple and yellow clusters. For example, the ellipsoid for position 146 illustrates that the C_α_ atoms in the green and red clusters have a much greater positional variation (the ellipsoides are bigger) compared to the cyan, purple and yellow clusters (Fig. 9). The ellipsoids show the variation of C_α_ coordinates of all structures belonging to each sub-master cluster. The sizes of the ellipsoids show that residue 146 of the green and red cluster’s structures varies a lot while the structures of cyan, purple and yellow cluster’s structures do not change as much.

Table 1 shows that datasets which belong to the green and red cluster have **a** = **b** unit cell values around 114 – 115 Å, while datasets of the other clusters have these values around 111 Å. This fact is shown in the cell parameter cluster in Fig. 4. The Table also reflects the fact that datasets belonging to the cyan, purple, and yellow clusters have higher resolution than those of the green and red clusters. The overall resolution of around 2 Å with good structure quality for each of the six structures is indicated by Rwork and Rfree about 20%.

When we align the model derived from the average cluster with the five major subclusters using FATCAT in rigid mode all the residues between 1 and 138 are well-aligned, but residues 139 to 146 increasingly diverge, as shown in Fig. 9.

### 3.4. Detecting dynamic behaviour via ligand-binding hot spots

FTMap shows six binding hotpots for each of the five structures. Among them, we observed the most different binding sites between red cluster’s structure (7KU2, cluster 140) and purple cluster’s structure (7KTZ, cluster 131). See Fig 10 for the differences for some of the six binding hotspots. Since these two structures belong to one each of the two different cell-based clusters, the differences provide strong evidence for the effectiveness of both cell-based and amplitude-based clustering in detecting polymorphs in the case of very small physical changes.

## 4. Discussion

Although both experimental work (Debrunner *et al*., 1982) and theoretical work (McCammon, 1984) established that dynamic behavior underlies most protein functions, crystallography was not regarded in the early years as an appropriate tool for investigating protein dynamics. An early review of protein crystallography concluded by stating that, “crystallographic methods are not suitable for the direct study of the dynamics of protein structure and interactions” (Stryer 1968). However, the presence of diffuse scatter implied that dynamic behavior was occurring within protein crystals (Caspar *et al.*, 1988), and crystal structures soon illustrated examples of protein dynamics (Ringe *et al*., 1985) induced by physical changes (for example temperature (Tilton *et al*., 1992), pH (Diao 2003), and ionic strength (Sanishvili *et al*., 1994)), and by chemical changes (for example adding denaturants (Dunbar *et al*., 1997) or ligands (Edwards *et al*., 1990)). However, Stryer’s 1968 assertion stands to this day in the sense that simple tools to identify dynamics from diffraction data are still not widely deployed, and consequently most crystallographic contributions to dynamics continue to be fortuitous. We propose here a method to suggest insights into the dynamics of proteins by systematic surveying of diffraction data for the presence of clusters. Once crystallographers are equipped with appropriate tools to identify clusters within aggregates of diffraction data, results indicating dynamic behavior may emerge routinely in many protein-crystallography projects.

A simple tool for identifying dynamic contributions in diffraction data must be as automated as possible, must present results in a way that is easy to interpret, and must be robust enough to identify small movements. The first of these requirements was simple to accommodate because we deployed our software within the existing KAMO software package, which we easily integrated into our existing version of the *fast_dp* automated data-analysis pipeline. We addressed the second requirement by incorporating visual tools such as systematic color annotation of clusters (Fig. 6), ellipsoid visualization for model variations (Fig 9.), and dot-plot visualization for data variation (Fig. 7).

**Figure 7.**
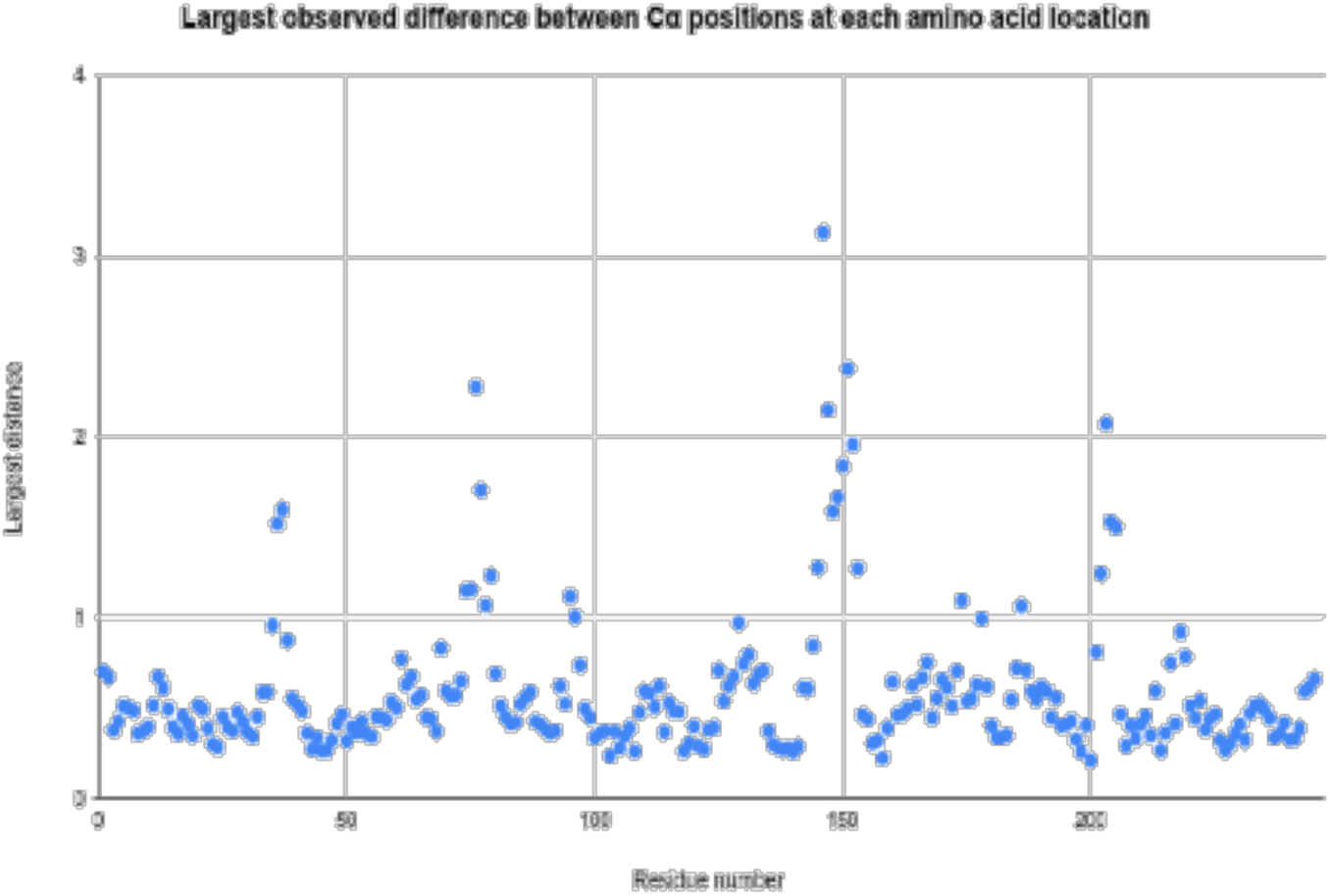
Largest observed difference between C_α_ positions at each amino acid location. To determine which regions of ChTg were most mobile in our data, we plotted the largest observed difference in the position of the C_α_ carbon for each of the 146 amino acids. The data illustrate one extended region with very large variation (between residues 146 and 151, in the vicinity of the missing loop). There are also two shorter regions with smaller variation around Ser75 and Val200.

**Figure 8.**
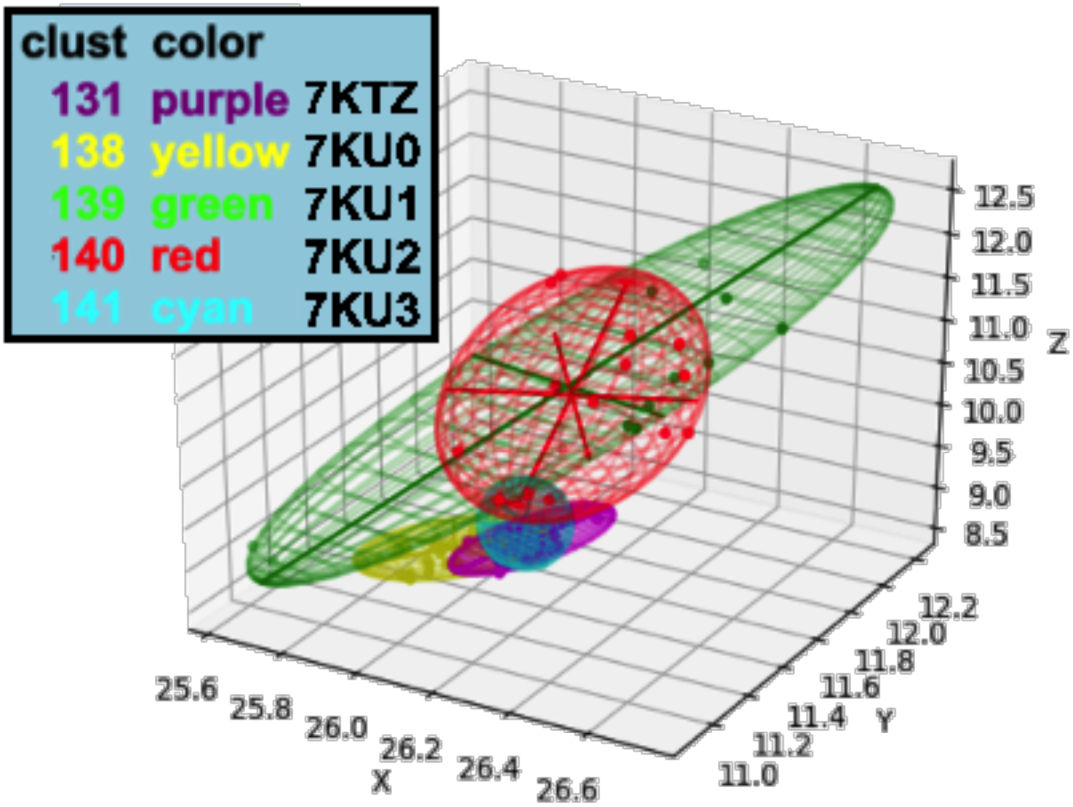
Using ellipsoids to illustrate variation in the C_α_ coordinate at position 146. We calculated five ellipsoids for each residue position, corresponding to the observed variation in the C_α_ positions at a specific residue for the green, red, cyan, purple and yellow data. The lengths of the perpendicular axes were determined using the minimum volume method (which minimizes the volume of the ellipsoid enclosing the data – see https://github.com/nsls-ii-mx/chymotrypsinogen and https://raw.githubusercontent.com/nsls-ii-mx/chymotrypsinogen/master/ellipsoid.py). This method optimizes the fit of each ellipse to the data, including the major axis in the direction of greatest variation. For example, at C_α_ position 146 (shown here) the green cluster yielded 18 structures with large variation in the [.2, −.8,0] direction. The volume of the ellipsoids indicates the overall variation in corresponding C_α_ positions. For example, at position 146 the green and red clusters yielded structures with much larger positional variation than the cyan, purple and yellow clusters.

**Figure 9.**
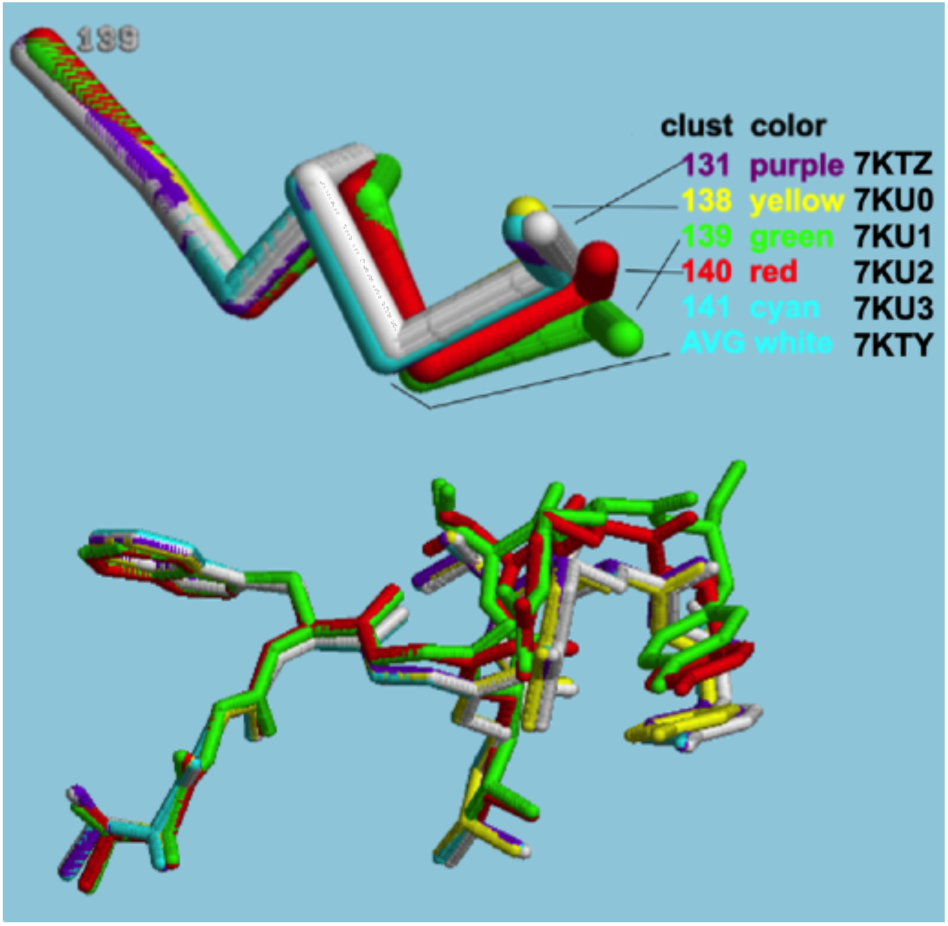
Variations in the clusters for residues 139 through 146. The overall average structure, 7KTY, which is cluster 145 in the dendrogram, is colored white. PDB entries 7KTZ, 7KU0, 7KU1, 7KU2, and 7KU3, which are clusters 131, 138, 139, 140, and 141, colored purple, yellow, green, red and cyan, respectively. The top half shows the variation in the backbone alone. The bottom half shows the variation with the side chains.

**Figure 10.**
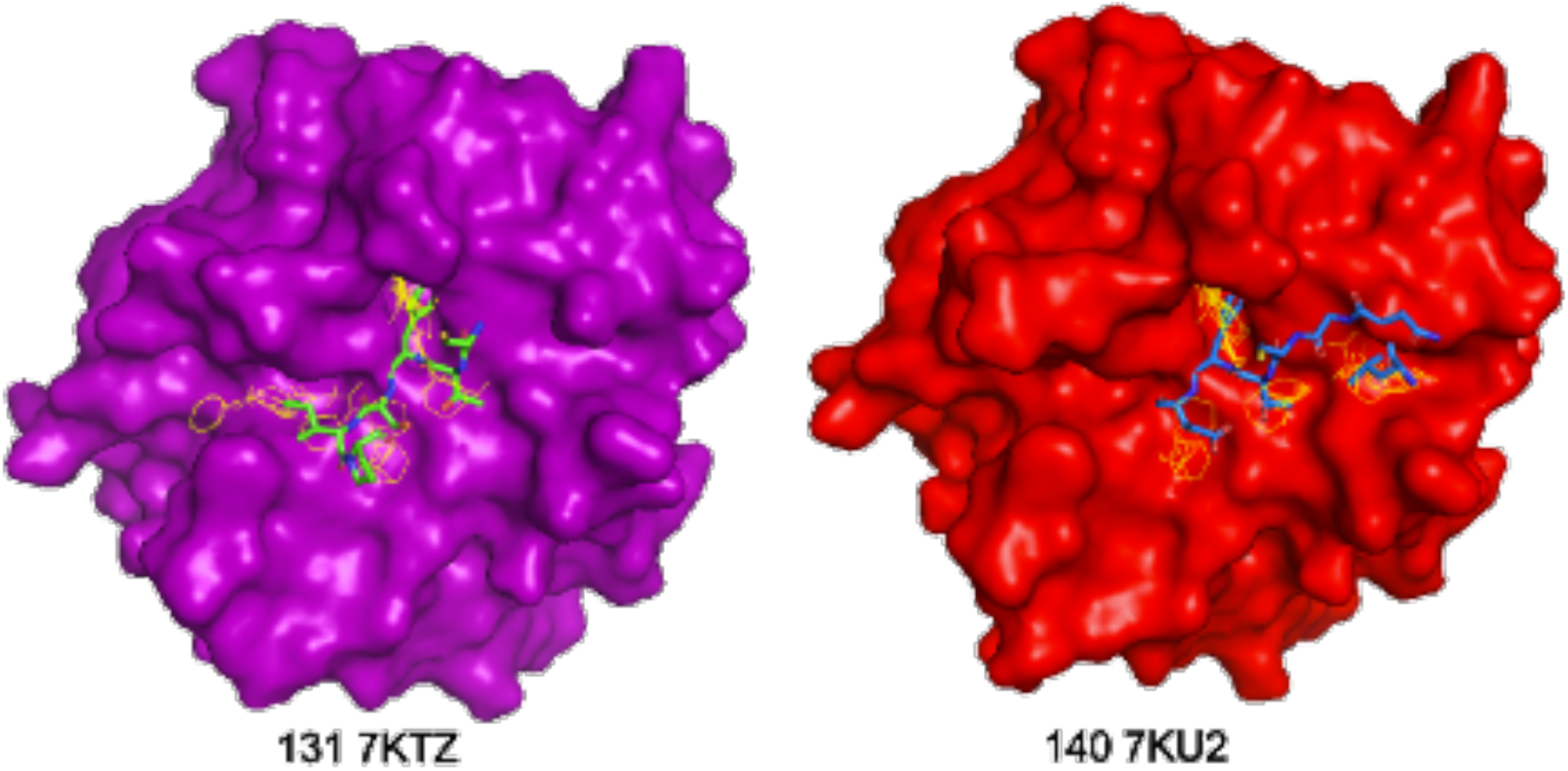
FTMap output comparing binding hotspots of clusters 131 (7KTZ) and 140 (7KU2). Figure 10 Left is an FTMap mapping result of model 131 (7KTZ, the purple cluster with **a**= 111.49 Â, as a Lee-Richards surface) overlapped with a wire-frame rendering of critical parts of Bowman-Birk protease inhibitor complex with chymotrypsinogen, PDB 3RU4 (Barbosa *et al.* 2007). Figure 10 Right is an FTMap mapping result of model 140 (7KU2, the red cluster with **a**= 115.57 Â, as a Lee-Richards surface) overlapped with a wire-frame rendering of critical parts of Kazal Type inhibitor, PDB 1CGI (Hecht *et al.* 1991).

The most difficult benchmark was the ability to differentiate clusters having limited dynamic contributions. We tested our techniques using our data from ChTg, which was not known at the outset to exhibit dynamic behavior. Many of the changes that we identified involved just a few amino acids. By combining the strengths of unit-cell clustering (ability to operate on highly granular data) and the strengths of diffraction-based clustering (sensitivity to very small structural changes), we believe that our technique will accurately identify relevant clusters of different structures hidden within highly similar data. Our method could detect different polystates with small coordinate differences, which in our case are less than 3Å in just two amino acids. In addition, the visualization tools that we created (color-based ellipsoid and scatter plots) allow easy identification of the highly-dynamic regions. This provides verification that our clusters are physically meaningful. These tools provide scientists a simple method to screen their data for dynamic behaviors.

High-data-rate crystallography represents a large and growing fraction of all crystallographic data. At synchrotrons, serial crystallography and combinatorial crystallography (*e.g.* fragment screening) produce robust streams of data from samples that are similar but not identical. Such data streams can be automatically clustered and be visually presented to scientists either to inform their main project or to yield serendipitous information that may expand their thinking of the system in question. XFEL light sources generate even larger data streams, with individual diffraction images that are derived nearly instantaneously from very small protein crystals. The great reduction in the time- and space-averaging in XFEL data (compared to synchrotron data) further increases the likelihood of obtaining data from crystals that are in different resolvable polystates. In either case, the data processing challenge is the same. One needs a data-clustering algorithm that is robust enough to work with mixed quality data, sensitive enough to partition all of the polystates that are present, and intuitive enough that investigators are able to identify useful clusters that represent biologically relevant polystates. Here, we present an algorithm that accomplishes all of these goals.

Our data processing and clustering are all automatic to reduce the time of screening and analyzing the molecules. We also do manual checking to verify that the automated processes achieved reasonable fits to density. However, it’s still a challenge for us if the data contain a lot of noise such as blur or unindexable spots. The problem may be solved by future research on spot finding and auto-indexing. In addition, we would like to test if different distance metrics could improve the accuracy of the clustering output and further improve the chances of detecting smaller potentially meaningful changes and to test if the tools could detect polystates well with datasets from other molecules so that we would have a comprehensive understanding about the efficiency of our clustering method.

## 5. Conclusions

Observing differences in protein structures, even small differences, could be meaningful and important. However, we usually miss the changes that are very small as they are very hard to measure. In this paper, we show how one might use the combination of our cell-based and structure-factor-based clustering methods to detect polystates of molecules. We applied these methods on ChTg data and were able to detect polystates with very small differences among five clusters of datasets. From these clusters, we built molecular structures and verified the differences among them. The combined method should help scientists to discover minor changes in molecules that are hardly noticeable by the change of cell parameters only.

To confirm polystate screening and verification, we propose using color-based visualization for distinct groups of datasets including dendrograms, ellipsoids, and scatter plots. The dendrogram shows the members of clusters with custom height cutoffs and the differences among those clusters. The ellipsoids show the variations of physical coordinates of clusters’ structures. And the scatter plot quickly shows cell-based clusters and their relations with structure factor-based clusters. Using the color-based plots, one could easily discriminate among groups of datasets. This visualization method is a fast way to screen a large number of datasets and point out which ones are important for further investigation.

## Acknowledgements

Data for this study were measured at beamlines 17-ID-1 (AMX) and 17-ID-2 (FMX) at Brookhaven National Laboratory’s National Synchrotron Light Source II (NSLS II).

Our thanks to Gregg Crichlow for careful and thoughtful review of both the PDB depositions and of this paper.

Our thanks to Frances C. Bernstein for many hours of copy-editing.

